# Epithelial convergent extension as a tuning process

**DOI:** 10.1101/2025.11.06.687029

**Authors:** Sadjad Arzash, Andrea J. Liu, M. Lisa Manning

## Abstract

Self-tuning—the ability of disordered systems to develop desired collective behaviors by tuning internal couplings in response to feedback—has recently emerged as a powerful framework for understanding adaptation in amorphous solids, mechanical metamaterials, and electrical networks. These systems can learn desired responses, encode memory, and robustly reorganize under repeated stimuli, much like artificial neural networks but without requiring processors to adjust their weights. Here, we extend this paradigm to morphogenesis and show that the epithelium can be viewed as tunable matter and that epithelial convergent extension (CE) can be understood as a self-tuning process. Using a vertex model with active interfacial tensions, we systematically compare distinct tension-update strategies, including externally imposed shear, global gradient descent optimization, and decentralized local feedback rules. We find that while all methods can generate tissue elongation, only local orientation- and length-sensitive rules reproduce key experimental features of CE, such as supracellular actomyosin pattern formation, cell shape changes, and junctional alignment. In contrast, global optimization produces homogeneous tension patterns and mechanically fragile states. By interpreting CE through the lens of tuning, our framework bridges the physics of tunable matter with developmental biology, revealing how simple, local rules enable tissues to efficiently orchestrate complex morphogenetic outcomes through decentralized mechanical adaptation.

## INTRODUCTION

Can we better understand the mechanisms that drive morphogenetic processes such as convergent extension (CE) by interpreting such processes as a form of *self-tuning* [1]? Embryonic tissues, like mechanically-tunable matter [2–4], can reorganize their internal mechanics to achieve targeted deformations. Here we introduce a framework that treats junctional tension remodeling as a self-tuning process and systematically compare global and local tension-adaptation strategies within a vertex model for confluent biological tissues [5, 6].

This framework allows us to ask and answer new types of questions: Do multiple different tuning processes perform equally well at generating convergent extension, or is there an optimal process that is vastly better than the others? Are some processes more robust than others? Are *global* strategies, which require processing information about all the junction tensions at once, substantially faster than local strategies, which only require each junction to process information about itself and/or nearest neighbors? Are there observable signatures that distinguish between local or global strategies, or even allow us to pinpoint a specific learning strategy that best matches experimental data?

Answering these questions not only provides a systematic method for testing and comparing specific mechanisms that have been proposed to drive morphogenetic processes, but also identifies a suite of possible (and sometimes novel) mechanisms that can be used to analyze robustness and the space of evolutionary possibilities.

Although the framework we develop here can be extended to other morphogenetic processes, we focus here on convergent extension in *Drosophila* because it is a stereotyped and very well-characterized global tissue deformation in a simple epithelium. Generally, convergent extension is a morphogenetic movement found across many different species in which epithelial tissues elongate along one axis while narrowing along the perpendicular axis, which resembles global pure shear when the overall area remains unchanged. Convergent extension underlies axis elongation in both invertebrates and vertebrates, including the *Drosophila* germband [7, 8]. At the cellular scale, CE is driven by polarized neighbor exchanges, cell shape changes, and directed cytoskeletal activity [7, 9].

A defining feature of epithelial CE is the anisotropic distribution of contractile forces at cell-cell interfaces, mediated by planar-polarized myosin II [10].

In *Drosophila*, myosin accumulates at anteriorposterior junctions, where it promotes directional contraction and intercalation, while being depleted from dorsal-ventral junctions [7, 10, 11]. These forces are dynamically remodeled and show feedback sensitivity to tension, with increased myosin localization at stressed junctions [11]. Moreover, supracellular actomyosin patterns such as cables and bridges [12] emerge along DV-aligned interfaces, suggesting that coordinated multicellular structures contribute to global tissue flow [8]. More recently, careful investigations have simulated and/or inferred specific rules for the molecular feedbacks that drive observed myosin and tension reorganization in epithelial tissues [12–28]. Given these myriad different proposed rules for feedbacks, it would be useful to have a systematic method for comparing them to one another and verifying how effective or robust they are at accomplishing developmentally-important tissue-scale tasks.

Theoretical models, particularly vertex models, have provided a natural language for testing whether local mechanical rules are sufficient to generate large-scale tissue deformation [9, 29]. Simulations have shown that anisotropic tensions—whether imposed externally or generated by molecular feedback—can drive T1 transitions and axis elongation. Based on experimental observations, recent work has introduced dynamic remodeling of edge tensions based on geometric or mechanical feedback [12– 14]. This work confirms that vertex models with several different classes of feedback rules can successfully generate convergent extension, and also match observed geometric features of tissues. Therefore, our goal is to compare the efficiency and robustness of some of these different rules or strategies in accomplishing the task of convergent extension.

To do so, we leverage recent advances in the physics of tunable matter. Proteins can be viewed as examples of tunable matter, in which the choices of one in 20 amino acids at each point in the sequence (the “tunable degrees of freedom”) determine protein structure and function. Proteins have been tuned by the biological evolution process. In contrast, there are self-tuning systems, as has been proposed for the actin cortex [30] where adaptation is understood as the systematic self-tuning of internal “tunable” degrees of freedom characterizing effective interactions, in response to feedback [1, 31–34]. In this picture, disordered networks learn by reorganizing their couplings – i.e. ‘tunable degrees of freedom’ – to reproduce desired responses encoded by a cost function, in direct analogy to artificial neural networks, where during training network weights are updated to minimize a loss function – the difference between the prediction and the correct answer for a training data set [35]. In neural networks and in computer optimization of materials [36, 37], one generally performs global optimization of the cost function with respect to the tunable degrees of freedom. However, in physical systems it is not always possible or desirable to have an on-board global processor, and instead one must use decentralized local update rules [1]. In many cases, these rules are inspired by contrastive learning approaches introduced in models for real, biophysical neural networks [31, 32, 38, 39]. Self-tuning physical systems appear to share important features with artificial neural networks, such as the fact that they generally find good solutions even though they are vastly overparameterized – i.e. they have many more tunable degrees of freedom than the number of features observed in the data [33].

The analogy to tissues is striking: like self-tuning physical networks, embryonic epithelia undergo targeted deformations by remodeling their internal mechanics in response to local signals. Indeed, we recently showed that tissues can fluidize by tuning cell-level properties such as target shapes and areas [40]. Here, the tension along each edge in a vertex model is treated as an individually tunable degree of freedom, and the learning task is defined by a cost function that specifies continuous elongation of the tissue along the anterior-posterior axis at fixed area. We compare diverse tension-adaptation processes, including global optimization of the cost function.

We find that several processes are quite efficient at generating robust convergent extension, reminiscent of overparameterization results for artificial neural networks [41]. Nevertheless, only processes based on length- and orientation-sensitive local rules can reproduce experimentally observed *in vivo* features such as supracellular myosin patterns and increasing average cell shape. Indeed, we find that a specific local rule inspired by contrastive learning best captures the experimental data. While global gradient descent is slightly faster at generating CE as the edge tensions are tuned, it produces qualitatively different solutions and tension patterns from those seen in experiments, indicating that experimental data can be used to distinguish whether local or global tuning processes are employed for a given task.

## METHODS

### Vertex model with active interfacial tensions

We model epithelial tissue dynamics using a modified version of the standard two-dimensional vertex model [5, 6, 42]. In this framework, each cell is represented as a polygon, and the degrees of freedom are the positions of the vertices shared between neighboring cells.

The mechanical energy of the tissue is given by

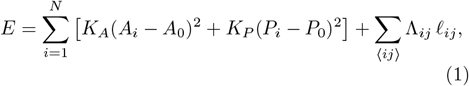

where the first summation runs over all cells, and the second over all edges between adjacent cells. Here, *A*_*i*_ and *P*_*i*_ are the area and perimeter of cell *i*, while *A*_0_ and *P*_0_ are their corresponding target values. The coefficients *K*_*A*_ and *K*_*P*_ control the stiffness associated with area and perimeter deformations, respectively. The third term describes interfacial tension: each cell-cell junction *ij* of length 𝓁_*ij*_ is associated with an active tension Λ_*ij*_, which may evolve dynamically. The inclusion of the tension term allows us to model active mechanical processes at cell interfaces, such as actomyosin contractility and tension remodeling.

Note that each edge also experiences a passive tension contribution, denoted by 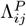, which arises from the perimeter elasticity of the adjacent cells. For an edge *ij* shared between cells *m* and *n*, this passive tension is given by 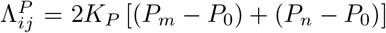. The total tension on edge *ij* therefore becomes

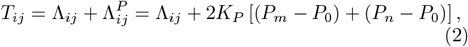

where Λ_*ij*_ is the active tension component.

We perform our simulations using the open-source `cellGPU` framework [43]. To initialize the tissue, we place *N* random seed points in a square periodic box of size 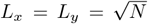 and construct the corresponding planar tissue using a Voronoi tessellation. Unless otherwise specified, we use parameters *K*_*A*_ = *K*_*P*_ = 1, with target area and perimeter values *A*_0_ = 1 and *P*_0_ = 3.7, which ensures that the passive contributions to the overall tension are small relative to the active tensions. We verified that our main results are robust to variations in *P*_0_, provided the tissue remains in a solid-like state, which requires *P*_0_ *<* 3.9. We also studied an active tension model [12, 44] where there are no passive tensions and the energy is linear in the edge lengths with no quadratic term. We verified that the results are also robust to this change in model (see SI). As highlighted previously [12, 13], the latter model can be difficult to regularize numerically, especially for larger systems, so we focus on vertex model results for the remainder of the main text.

To prepare the initial configuration, we first minimize the mechanical energy without the active tension term in to obtain a force-balanced state. We then initialize the active edge tensions Λ_*ij*_ to match the corresponding passive tensions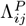 and perform a second minimization of the full energy, including the active contribution, to obtain the final initial state. To ensure that the initial simulated tissues matched experimental conditions, we tuned the distribution of active edge tensions so that the average cell shape index and the fraction of vertically oriented edges in the relaxed state matched those measured in *Drosophila* epithelia (Fig. 3). Specifically, we iteratively varied the mean and standard deviation of active tensions, minimized the energy, and compared the resulting tissue properties to experiment until convergence was achieved. For all our simulations, T1 events are performed when an edge length becomes shorter than a cutoff. After T1, we also reset the active edge tension Λ_*ij*_ to its average value ⟨Λ⟩ (see SI for more details).

### Different methods for achieving a convergent-extensional flow

To investigate how epithelial tissues undergo convergent extension, we implemented several distinct methods for driving tissue elongation.

#### External shear deformation

One straightforward approach to simulate convergent extension is to impose a global, externally applied pure shear deformation. In this method, we deform the entire tissue, along with the periodic boundary vectors, using an affine transformation

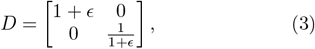

where *ϵ* is the imposed strain. This deformation stretches the tissue along the *y*−axis while compressing it along *x* − axis, mimicking the global shape changes observed during convergent extension in vivo.

After each incremental deformation step, we minimize the tissue’s mechanical energy at fixed strain to obtain a force-balanced configuration. While the active edge tensions Λ_*ij*_ remain fixed at their initial values throughout this protocol, the total edge tensions *T*_*ij*_ evolve due to changes in cell perimeters via the perimeter elasticity term. This stepwise procedure allows us to systematically investigate how cell shapes, junctional orientations, and tension distributions respond to increasing strain. Although this approach does not incorporate active cellular processes, it serves as a valuable baseline for understanding the mechanical response of passive tissues to externally imposed anisotropic deformations. This rule is also biologically plausible, as the germband in *Drosophila* is surrounded by other tissue types undergoing different morphogenetic transformations, which have been suggested to apply external shear to the germband [45].

#### Global gradient descent on edge tension degrees of freedom

An alternative to externally imposed deformation is to treat the active edge tensions Λ_*ij*_ as tunable degrees of freedom that actively drive tissue remodeling. In this approach, we induce convergent extension by dynamically adjusting the edge tensions via gradient descent on a global cost function. Here, the ‘task’ we are investigating is how effectively the tissue performs global convergent extension. Since the *Drosophila* tissue roughly maintains the same area during the process, we approximate convergent extension as global pure shear. Therefore, we define the cost function as

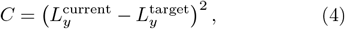

Where 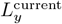 is the current length of the tissue in *y*-axis, and 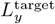 is a prescribed target length that represents the desired deformation state. This function could be alternatively defined in terms of *L*_*x*_. The evolution of edge tensions follows a gradient descent on this cost function

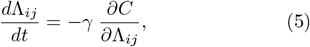

where *γ* is the tuning rate that controls how rapidly tensions are updated.

Note that this definition does not encode a stopping point for the CE process – under this definition of task the tissue will elongate indefinitely. Recent work suggests that additional processes or tissue features may lead to arrest of tissue elongation [12], and such tasks could be incorporated into a time-varying cost functions [46, 47] in future work.

At each tuning step, after updating the tensions, we minimize the mechanical energy of the tissue with respect to the physical degrees of freedom (vertex positions) and the global pure shear strain *ϵ*, which parametrizes the deformation of the simulation box. The box deformation is updated by minimizing the total energy with respect to *ϵ*, i.e., by solving *∂E/∂ϵ* = 0 (see SI for more details). Clearly, this rule is not biologically plausible, as it would require an external processor that knows the state of all edges simultaneously and uses that information to compute a global gradient. However, it is by definition the optimal (fastest) path for achieving global pure shear via altering edge tensions, and so it provides a useful comparison for other strategies.

#### Internally-driven deformation via an experimentally-derived local tension rule

Beyond external shear or global optimization, we are interested in whether there are local rules – rules that depend only on features of the edge itself and/or its neighbors – that could drive the tissue to converge and extend. Such rules could be biologically plausible, as they do not require an external processor, and in fact, most of the previously proposed adaptive rules for edge tensions [12–14, 44] are local. A challenge is that many such local rules, and similar ones developed for other confluent tissues [48, 49], generate temporal dynamics for myosin interacting with other spatial fields, which become dynamically complex and difficult to simulate/study [50].

For simplicity, we focus here on local rules that do not possess any internal dynamics. In some cases, they can be understood as the limit of dynamical models where molecular feedback processes are fast compared to cellular rearrangement timescales. One local rule is inspired by a simple experimental observation – that myosin accumulates on edges oriented along the DV axis. As shown in Fig. 1(c), the observed pattern is well approximated by an equation where the concentration of myosin *c*_*myo*_ is related to the angle that the junction makes with the AP axis *θ*_*ij*_:

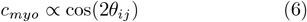

**FIG. 1.**
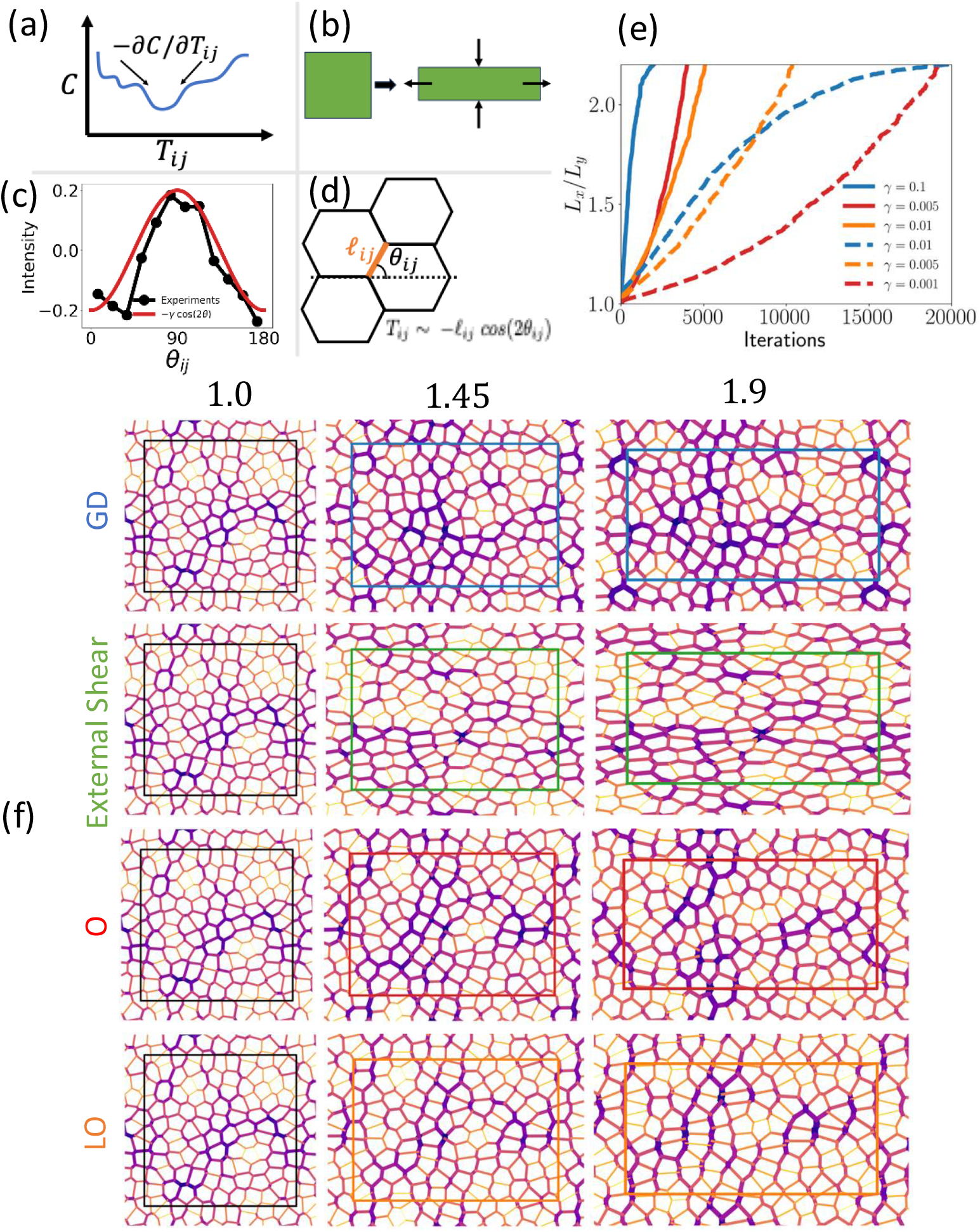
(a) Schematic of the global gradient descent rule. (b) Schematic of the external pure shear deformation rule. (c) Myosin intensity versus cell–cell edge orientation; the black curve shows experimental data adapted from [8], and the red curve shows the fit − *γ* cos 2*θ*_*ij*_. (d) Schematic of the local tension rule on cell edges, 𝓁_*ij*_ cos 2*θ*_*ij*_. (e) Tissue aspect ratio *L*_*x*_*/L*_*y*_ versus the number of edge tension update iterations for three remodeling strategies: the local rule 𝓁 cos(2*θ*) (orange), the global gradient descent rule (blue), and the orientation-only rule cos(2*θ*) (red). Each method is shown for different tuning rates *γ* as indicated in the legend, with data averaged over multiple samples. All three methods are capable of driving tissue elongation to arbitrary aspect ratios, with the number of iterations required depending on tuning rate. (f) We show representative tissue snapshots illustrating convergent extension under different remodeling protocols. Each row corresponds to a different mechanism, with columns showing configurations at increasing aspect ratios: the initial state at *L*_*x*_*/L*_*y*_ = 1.0, followed by intermediate (1.45) and elongated (1.9) states. Edge thickness is proportional to total tension, and colors indicate tension magnitude (darker means higher tension). Notably, at high aspect ratio, the local rule produces cable- and bridge-like structures of aligned high-tension edges [12], a feature absent in both the GD and externally sheared tissues.

This suggests an orientation-dependent remodeling rule, under the assumption that the active tension is proportional to the concentration of myosin, and that tensions follow this rule along their dynamic trajectory. Specifically, Wang et al. [51] investigated a phenomenological rule that evolves edge tensions based solely on their orientation

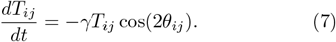

To implement this rule, we update the total edge tensions *T*_*ij*_ at each time step according to the specified local feedback mechanism. We then minimize the mechanical energy defined in (1) with respect to both the vertex positions and the global pure shear degree of freedom. The tuning rate and time step are chosen to ensure numerical stability (see SI for details).

We note that the noisy data from Fig 1(c) doesn’t strongly constrain the functional form for the dynamical rule; therefore we have also studied dynamical rules where the update is proportional to a triangular or step function in *θ* and find that the results are quite similar to that for the *cos*(2*θ*) case. (See SI for details). As this local rule is inferred from data, it is clearly biologically plausible. From here on, we refer to this orientation-dependent local rule as the local rule “O”.

#### Internally-driven deformation via a theoretically-derived local tension rule

Another goal of this work is to understand whether interpreting morphogenetic processes as tuning rules can help generate new, testable hypotheses for underlying mechanisms that drive such processes. In the field of tunable matter, an effective local rule is “directing aging,” in which the desired state is applied as a boundary condition and the energy of the physical system under that boundary condition is minimized with respect to the tunable degrees of freedom [52–54].

Inspired by these ideas, we seek a local rule that lowers the energy in the direction of a pure global shear deformation. To ensure the rule is local, we study how each edge should change itself in order to lower the energy in the direction of pure shear. In the total energy expression of (1), the third term represents the total tension energy *E*_Λ_, which depends on both the edge lengths 𝓁_*ij*_ and the interfacial tensions Λ_*ij*_. We study how the linearized version of global pure shear deformation (3),

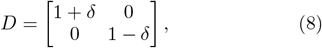

impacts the energetic cost, governed by how *E*_Λ_ changes with *δ*. Specifically, the derivative

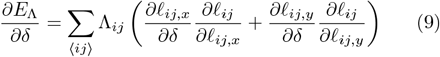

captures how edge tensions contribute to the resistance against shear. The expression inside the parentheses simplifies using *∂*𝓁_*ij,x*_*/∂δ* = 𝓁_*ij,x*_ and *∂*𝓁_*ij,y*_*/∂δ* = − 𝓁_*ij,y*_. Substituting these, we obtain

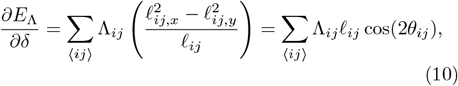

where *θ*_*ij*_ is the angle of the edge with respect to the *x* − axis. To reduce this resistance locally—i.e., to lower the energy of the pure shear mode—we propose a feedback rule that modulates the total tension *T*_*ij*_ based on each edge’s length and orientation. This local rule is given by

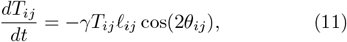

where *γ* is the tuning rate. This rule decreases tension along edges aligned with the *x*− axis (AP axis of embryo), where cos(2*θ*) *>* 0, and increases tension along the *y* − axis (DV axis of embryo), where cos(2*θ*) *<* 0. As a result, the tissue becomes more compliant in the direction of convergent extension and more resistant in the orthogonal direction, biasing the tissue toward an elongation response. This rule is implemented using a similar numerical framework as the rule in the previous section, see SI for details.

This rule is conceptually similar to the *directed aging* protocols used to train disordered materials [52, 53]. However, unlike conventional directed aging, where the training arises from repeated boundary-driven loading, the present local rule represents a decentralized feedback mechanism that modulates junctional tensions based on local geometry. This rule is simple, local, and orientation-based – making it biologically plausible. From a theoretical perspective, because it involves a single-edge based approximation to the cost function, it is the best that the tissue can do at generating pure shear if each edge has information only about itself.

Comparing to the simple experimentally-based rule from the previous section, we find that an additional length-dependent factor 𝓁_*ij*_ appears in the feedback loop. As shown below, this multiplicative length factor is not merely a refinement – it is essential for accurately reproducing key experimental observations. In particular, it generates cell shape changes that are better match experimental data, and suggests that effective tension remodeling *in vivo* may be sensitive not only to edge orientation but also to edge length. From here on, we refer to this length- and orientation-dependent local rule as the local rule “LO”.

## RESULTS

### Distinct convergent-extension flows under global versus local tension dynamics

Figure 1 compares the time evolution of the tissue aspect ratio *L*_*x*_*/L*_*y*_ under the global gradient descent (GD) method, schematically illustrated in Fig. 1(a) and the two local tension remodeling rules described earlier (panels c,d). We also show results for externally-imposed shear, which is not a tuning process (panel b). As shown in the top panel, all methods are capable of driving tissue elongation to a desired aspect ratio. The number of iterations required to reach a given aspect ratio depends on the tuning rate *γ*: larger values of *γ* lead to faster elongation in both methods. However, excessively large *γ* values can lead to numerical instabilities.

We note that for all three tuning processes there are many possible tissue states that the system can reach at the end of each tuning step, specified by the values of the tunable degrees of freedom (active tensions) and physical degrees of freedom (vertex positions). The aspect ratio curves shown here represent quantities averaged over ensembles of the tissue states that are reached at each step by each of the processes, each starting with a different initial state of the tissue. For a given initial condition and tuning process, the dynamics exhibit alternating regimes of smooth elastic deformation and discrete plastic events associated with T1 transitions [55], which appear as sudden jumps in aspect ratio.

Fig. 1 (f), we show representative tissue snapshots at three stages of deformation for each method. The left column shows the identical initial tissue configuration used for all methods. As the tissue elongates, distinct features emerge. The local rule LO produces prominent, extended, linear structures of high-tension (thick and dark) edges, reminiscent of high myosin patterns that are aligned preferentially along the *y*-axis (corresponding to the dorsal–ventral direction in embryos). These supracellular structures span multiple cells. In contrast, the GD method maintains a more homogeneous tension distribution and lacks such extended linear structures. Notably, similar actomyosin patterns have been observed in vivo during convergent extension (e.g., in Drosophila germband extension [9, 11] and vertebrate neural tube closure [56]). Recent work suggests that such extended tension patterns can be categorized into two classes, *tension cables* and *tension bridges*, based on the geometric properties of the underlying tension triangulation [12, 13], and analyzing these features in tuning processes would be an interesting direction for future work.

The external shear method, while effective at generating tissue elongation, yields tension patterns that are markedly different, including elevated tensions along the *x*-axis, further emphasizing the distinct mechanical signatures of each protocol.

### Comparing tuning processes: misalignment with gradient descent and other quantitative measures

A useful outcome of interpreting morphogenetic events as tuning processes is that we can precisely quantify how well or efficiently different processes perform in executing a specified task over a specified set of tuning degrees of freedom.

The most straightforward method to compute how efficiently degrees of freedom are tuned to execute a specific task is to study how quickly a process moves downhill in the landscape of the cost function defining the task. In our case, since the gradient descent rule is by definition the optimal strategy for moving downhill, we define an *optimization misalignment* that captures how far local updates deviate from the globally optimal direction provided by GD. Let 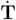 (*ϵ*) denote the instantaneous tension-update vector (stacking all *T*_*ij*_ updates) at strain *ϵ*. We compute a pointwise directional discrepancy via the dotProduct

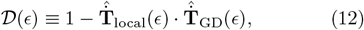

where hats denote normalization. The *cumulative* misalignment is then

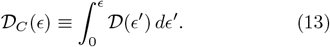

By construction, GD has *𝒟*_*C*_(*ϵ*) = 0, whereas local rules accumulate misalignment as strain increases (Fig. 2a), confirming that they travel a suboptimal path in cost space.

**FIG. 2.**
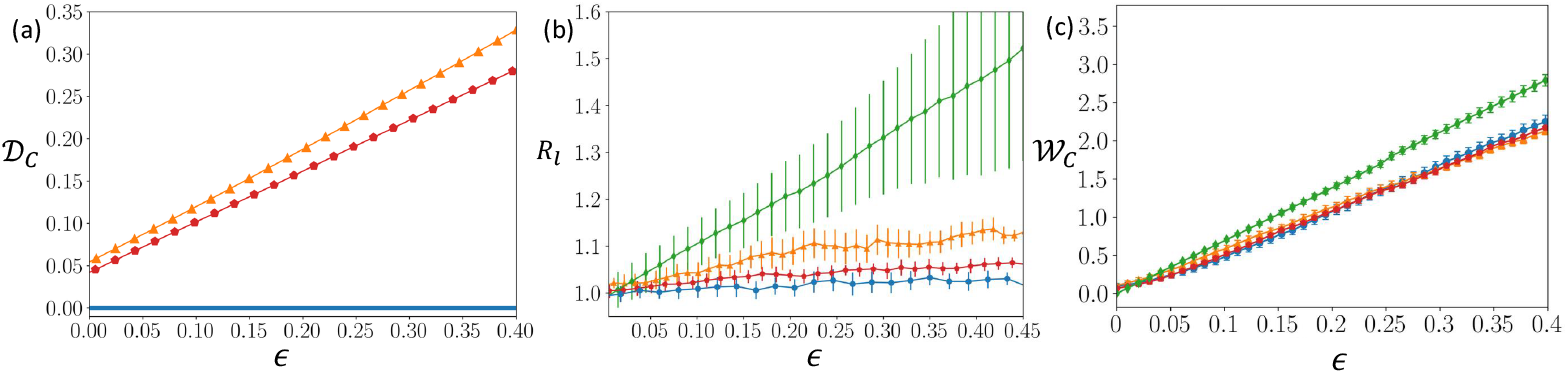
Comparison of properties for different tuning processes during convergent extension. (a) Cumulative optimization misalignment *𝒟*_*C*_ as a function of strain for different methods. This measure captures the deviation of local feedback strategies from the globally optimal update direction. Increasing misalignment with strain highlights the mechanical “cost” of decentralized adaptation relative to global optimization. (b) Ratio of projected edge lengths in the *x*-direction to those in the *y*-direction, *R*_𝓁_, as a function of strain. This metric quantifies anisotropy in edge orientations, distinct from tissue aspect ratio. The global gradient descent (GD) method yields the lowest anisotropy, consistent with more uniform edge distributions, while external shear produces the largest *R*_𝓁_; local rules fall between these extremes. (c) Cumulative work performed per unit area *𝒲*_*C*_ as a function of strain. This metric quantifies the total energetic cost required to deform the tissue under each remodeling protocol. External shear requires the largest cumulative work, while gradient descent (GD) and local feedback rules spend less work overall. **Symbols:** blue circles = GD; orange triangles = local rule LO; red pentagons = local rule O; green diamonds = external shear.

The cumulative misalignment *𝒟*_*C*_(*ϵ*) is roughly linear in *ϵ*, indicating that the differential misalignment is *𝒟* (*ϵ*) is independent of strain. The slope is similar for the two local rules studied, though somewhat surprisingly, the local rule O is slightly more efficient by this metric. Unfortunately, it is not obvious how to compute _*C*_(*ϵ*) from experimental data. On the other hand, it is clear that imperfect projection of a local rule onto the direction of steepest descent does not compromise the system’s ability to perform a task, consistent with observations in other systems [32, 57].

A second method to quantify efficiency, which can also be computed for experimental data, is inspired by the triangle method for quantifying contributions to global deformation developed by Merkel et al. [58]. In that work, the authors demonstrate that global pure shear can be written as a sum of five components related to the geometry of a celullar packing. Two of those components, cell extrusion and cell division, do not occur in our system, and a third, due to correlation effects, is generally found to be small in both simulations and experiments of Drosophila epithelial tissues [58]. The remaining two terms correspond to cell neighbor rearrangements and changes to the elongation of individual cells.

We postulate that there might be a difference in how different tuning processes distribute global shear between these two terms – some may generate more cellular rearrangements, while others may incur a geometric cost by increasing cell elongation along the direction of shear. To quantify this, we measure a purely *geometric* cost defined as

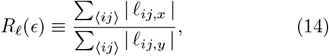

the ratio of projected edge lengths along the *x* and *y* directions at fixed tissue strain *ϵ*. Unlike the tissue aspect ratio, which is set by the strain protocol, *R*_𝓁_ reports how each rule redistributes junction orientations and lengths within that global constraint.

A comparison of *R*_𝓁_ for different tuning processes is shown in Fig. 2b. Consistent with intuition, externally imposed shear yields the largest *R*_𝓁_ because the boundary deformation directly elongates cells in the *x*-direction (Fig. 2b). In contrast, the global gradient descent (GD) method produces the lowest anisotropy, indicating a more symmetric, uniform allocation of edge orientations. The local rules—which rely only on decentralized, geometry-sensitive updates—fall between these extremes.

The previous metrics focused on the cost function associated with the task. However, because the tissue is a physical system, there is also a separate energetic cost to any process, associated with the work required to accomplish it. Therefore, we compute the cumulative work per unit area associated with the each local rule as a function of the applied strain *ϵ*:

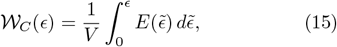

As shown in Fig. 2c, this work is nearly identical for both the global and local rules, with the local rules requiring slightly less work over the largest strains. Given that the paths and cell geometries generated by the different rules are quite distinct, the similarity in the work is quite striking, and suggests that many different rules are able to find “good solutions” from an energy cost perspective.

### Local rule LO best captures cell-shape changes

Finally, we ask whether there are experimental observables that might distinguish between different learning rules, allowing us to infer which rules might be operating in a given *in vivo* system. Here, we focus on experimental quantities that have already been reported elsewhere in the literature.

One geometric property that has been reported is the statistics of edge orientations. Specifically, Ref. [11] measured the fraction *f*_*v*_ of edges whose orientation lies within ±15^*°*^ of the *y*-axis. Here, we adapt their reported data to plot it as a function of tissue strain, shown by the black dots in Figure 3a. This data exhibit a clear increase in D–V alignment with increasing strain: as tissues elongate, cells not only stretch but also reorient their junctions to align perpendicular to the extension axis.

We can compute this exact quantity for each of our learning rules as well. The external shear case (green) incorrectly predicts a decrease in *f*_*v*_, since passive stretching tends to rotate existing edges toward the extension direction. Similarly, the GD method and the rule O (blue and red curves) fail to produce significant alignment. Only the feedback rule LO (orange) captures the experimentally observed trend, suggesting that effective tension remodeling must incorporate both edge orientation and length to selectively stabilize D–V junctions. However, none of the rules studied here quantitatively captures the black experimental data, where the fraction of vertical edges increases extremely quickly and then levels off. Since this metric relies on an arbitrary cut-off on what counts as vertical, which induces significant noise, it would be interesting to revisit these experiments to look at the statistics of all orientations [8] as a continuous function of strain.

Another quantity that has been measured in experiments is the mean cell shape index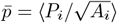, which increases with increasing cell elongation. This quantify was measured in *Drosophila* CE by Wang et. al [51], and in Figure 3b we have replotted that data as a function of strain (black circles).

**FIG. 3.**
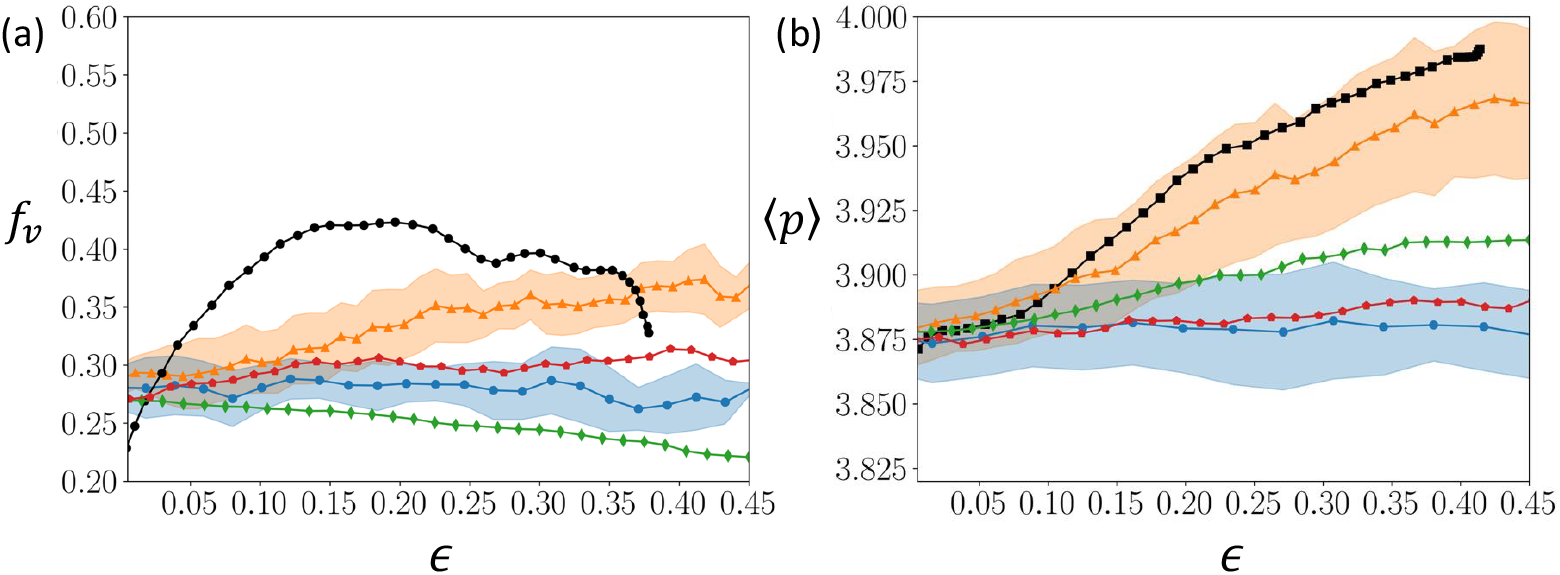
Macroscopic mechanical observables under different convergent-extension mechanisms. (a) Fraction of vertically oriented edges, *f*_*v*_, defined as the fraction of edges with orientation angle *θ* satisfying 75^*°*^ *< θ <* 105^*°*^, where *θ* is measured with respect to the horizontal (*x*-) axis. Experimental data (black circles) are replotted from Ref. [11]. Simulation results are shown for four protocols: external shear (green diamonds), global gradient descent (blue circles), local rule O (red pentagons), and local rule LO (orange triangles). Data represent averages over 50 independent samples; shaded regions denote standard deviations for the GD and LO rules. (b) Same as (a), but for the average cell shape index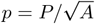, where *P* and *A* are the cell perimeter and area, respectively. Experimental data (black circles) are reproduced from Ref. [51].

Figure 3b compares the mean cell shape index,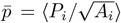, as a function of the overall tissue deformation for all four protocols—global gradient descent (blue circles), the heuristic orientation-only rule of Wang et al. (red pentagons), our directed aging rule (orange triangles), and pure external shear (green diamonds). Both gradient descent and the orientation-only rule severely under-predict the increase in cell shape with tissue deformation: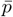 remains essentially constant, in stark contrast to experiments where cells progressively elongate. By contrast, the new rule reproduces the growth of 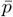 with strain, consistent with the experimental curve. Interestingly, externally imposed shear also drives cell elongation in a qualitatively correct trend, but in the absence of any active tension remodeling. This suggests that externally applied shear deformation alone can produce the right mean shape but cannot account for the underlying junctional mechanics.

Taken together, Figure 3 demonstrates that purely kinematic deformation or global optimization cannot fully recapitulate the coupled cell-shape and junction-alignment dynamics observed in convergent extension, but the local rule LO depending on both length and orientation performs substantially better. This is the main result of our study.

### Statistics of cell-scale features under different tuning processes

From the tissue states obtained via different tuning processes, we can quantify the statistical properties of various cell-scale features imprinted in the tissue, which can help us understand dynamical flows in the phase space of tunable degrees of freedom and quantify robustness.

We begin by examining the distribution of total edge tensions as tissues elongate, as we speculate that high tension edges are expensive and therefore not preferred. Fig. 4a) illustrates the strain evolution of the edges tensions normalized by their average value at zero strain, *T*_*ij*_*/* ⟨*T*_*ij*_⟩ _0_ under the local rule LO. As deformation progresses, we observe a gradual emergence of high-tension edges, manifested as a growing tail in the distribution. Concurrently, the curves shift to the right as the overall average tension in the tissue increases, reflecting the cumulative buildup of contractile forces over time. On the other hand, tissues undergoing convergent-extension via GD gradually decrease the high tension tails and do not exhibit a build-up of tension (Fig. 4d).

**FIG. 4.**
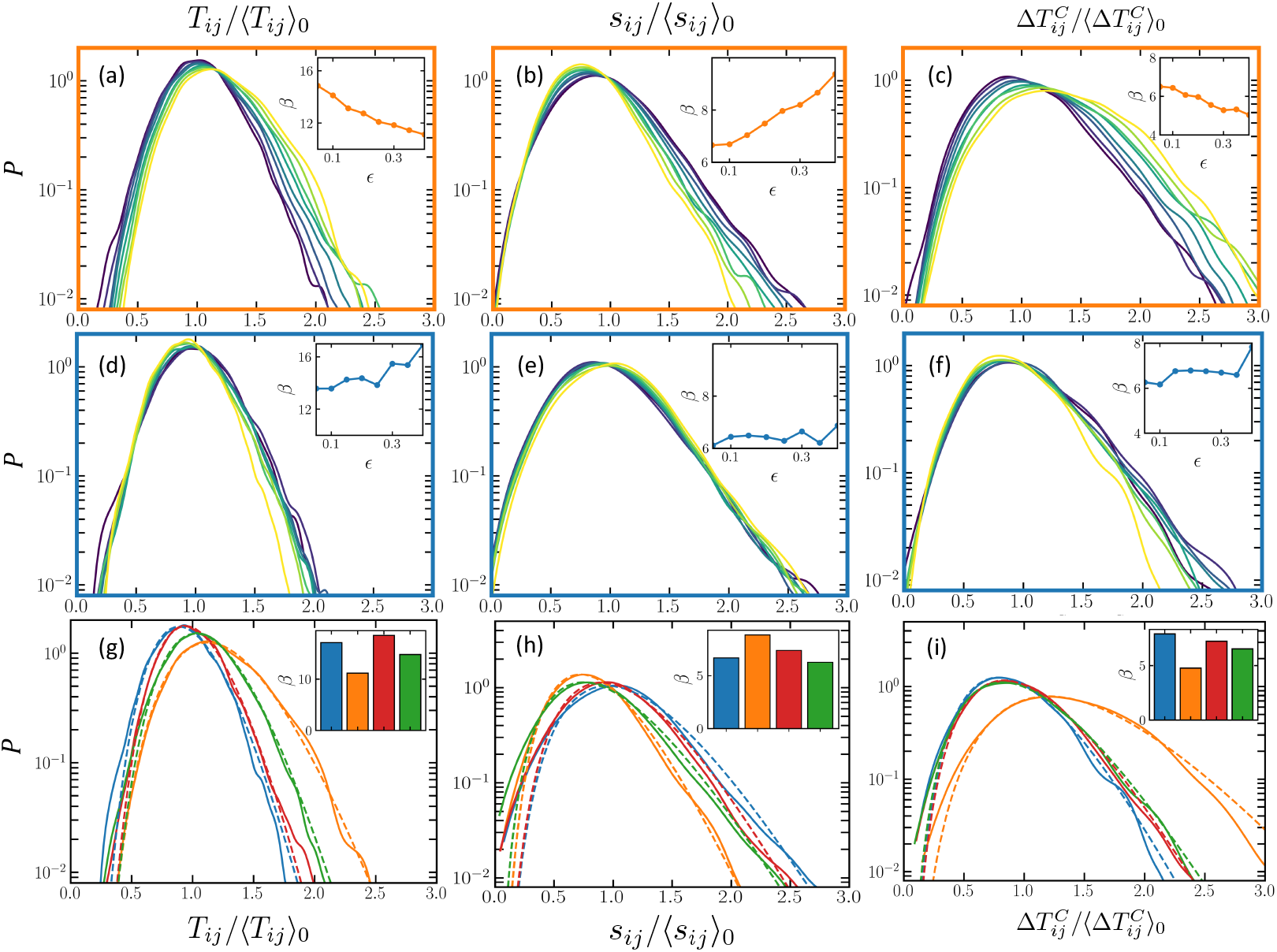
Probability distributions of microscopic mechanical observables under different convergent-extension mechanisms. (a–c) Probability densities of (a) total edge tensions *T*_*ij*_, (b) edge susceptibilities *s*_*ij*_, and (c) cellular tension anisotropy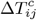for tissues undergoing convergent extension via the local rule *T*_*ij*_ ∝ 𝓁_*ij*_ cos(2*θ*_*ij*_). Colors indicate strain values increasing from *ϵ* = 0.05 to *ϵ* = 0.40 in steps of 0.05, with lighter shades corresponding to larger strains. (d–f) Corresponding probability densities for the same observables in tissues evolving under the global gradient descent (GD) method, with the same color scheme for strain. (g–i) Direct comparison of probability distributions at *ϵ* = 0.40 ±0.02 across different protocols: global GD (blue), local rule cos(2*θ*) (red), external shear (green), and local rule 𝓁 cos(2*θ*) (orange). Each distribution is aggregated over 50 independent samples. Together, these panels illustrate how microscopic mechanical statistics evolve during tissue deformation and how they differ between convergent-extension mechanisms.

Notably, the high tail of active tensions is significantly enhanced for the rule LO. To further quantify the behavior of the tails, we fit these distributions to a gamma function *f* (*x*) ∝ *x*^*α*−1^*e*^−*βx*^, where *α >* 1 characterizes the low tension onset, and *β* characterizes the fall off of the high tension tails (see SI). *β* decreases substantially as a function of the strain for the local rule LO, quanti-fying how the tails become heavier (Fig. 4a) inset), while under GD *β* increases, indicating the tails become lighter (Fig. 4d inset).

Next, we compare this distribution across different tuning processes and external shear conditions near the end of the process, at *ϵ* = 0.4. Fig. 4g compares the distributions of active tensions for all of the tuning processes, plus external shear, at a high tissue strain of *ϵ* = 0.4.

The local rule LO, which agrees best with experimental data, has a significantly enhanced high tail. In other words, high junctional tensions do not appear to be penalized, suggesting that active tensions costs are not an important biological constraint. The dashed lines in Fig. 4g show fits to a gamma distribution (see SI). The inset of Fig. 4g confirms that the LO rule yields a smaller *β*, consistent with a heavier-tailed distribution.

We next examine how susceptible edge tensions are to mechanical perturbations, as we speculate that highly susceptible edges make the system fragile with respect to mechanical perturbations, and may be penalized. To do so, we utilize the concept of *edge susceptibility*, following the approach by Guzmán et al. [59]. Edge susceptibility quantifies how sensitively the length of a given edge changes in response to infinitesimal forces applied at tissue vertices. It provides a local measure of mechanical responsiveness, capturing how edge geometry couples to global force distributions. Formally, for each edge *ij*, we define a susceptibility vector 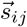, whose components describe how the length 𝓁_*ij*_ of edge *ij* changes in response to a unit force applied at each vertex *k* in the tissue. This definition reflects the linear response of edge *ij* to perturbations of *physical degrees of freedom* (vertex positions). The details of computation of this quantity are described in the SI. To summarize this information in a scalar form, we compute the squared Euclidean norm of the susceptibility vector 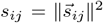. This scalar measure *s*_*ij*_ quantifies the *overall mechanical sensitivity* of edge *ij*, with larger values indicating edges that undergo significant length changes under force perturbations.

Strikingly, the high tail of the distribution of susceptibilities is suppressed for the local rule LO (Fig. 4b) compared to gradient descent ((Fig. 4e). Furthermore, as deformation progresses, we observe further suppression of high-susceptibility edges, manifested as a shrinking tail in the distribution. For gradient descent, on the other hand, we observe that the distribution of *s*_*ij*_ remains constant as tissue deforms (see Fig 4e). To quantify this evolution, we fit these distributions to a gamma distribution, as well. As shown in the insets to Fig. 4b) and e), *β* increases systematically under the local rule LO, reflecting the progressive suppression of highly susceptible edges, whereas it remains nearly constant for gradient descent. Again, we compare edge susceptibility distributions for all three tuning processes as well as external shear in Fig. 4h at the tissue strain of *ϵ* = 0.4. Clearly, the local rule LO (orange) has the most suppressed high tail of susceptibilities. This is further supported by the largest fitted *β* value (inset of Fig. 4h), consistent with the strongest suppression of high-susceptibility edges.

These results suggest that maintaining tissue configurations that are *mechanically more robust* may be biologically important.

We are also interested in tension heterogeneity within individual cells, as we speculate that it may be difficult for cells to maintain a large differential in surface energies between nearby surfaces. Therefore, we quantify the *cellular tension anisotropy*, defined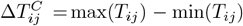, which measures the difference between the highest and lowest total edge tensions in each individual cell. For the local rule LO, we observe that the tail of 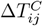 increases steadily with strain, indicating that local remodeling leads to growing cellular tension anisotropy during convergent extension (Fig. 4c). For gradient descent, on the other hand, this tail shifts towards lower values as the tissue elongates (see Fig. 4f).

As with the tension distributions, we are able to fit these data well with a gamma distribution and study how the high-tail parameter *β* changes with strain, shown in the insets to Fig. 4c and f. Fig. 4i shows that tension anisotropy is highest for the local rule LO. This suggests that maintaining low tension-anisotropy may not be particularly important biologically.

## Discussion

In this work, we reinterpret convergent extension (CE) in epithelial tissues as a tuning process, wherein junctional tensions are adaptively remodeled to achieve global tissue elongation. By comparing four distinct driving mechanisms—externally imposed shear, global gradient descent (GD), the length- and orientation-dependent local rule LO and an orientation-only local rule O, we show that only local rule LO robustly reproduces key experimental signatures of CE, including the formation of supracellular extended linear structures reminiscent of myosin patterns, progressive cell elongation, and junctional reorientation.

Our results highlight the biological plausibility of decentralized tension adaptation. The best-performing local rule,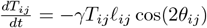, leverages purely local geometric cues to regulate edge tensions. Unlike global methods that require tissue-wide information flow, this rule achieves directional specificity and progressive remodeling through local computations alone. The inclusion of the edge length term 𝓁_*ij*_ amplifies feedback on longer junctions, leading to patterns of cell shape anisotropy and tension alignment that are most similar to experimental observations. Note that Fig. 2(a) shows that the LO rule (orange) has a larger cumulative optimization misalignment with the direction of gradient descent than the O rule, suggesting that the magnitude of the misalignment is not important as long as it is not too high.

Viewing CE through the lens of tuning also reveals possible tradeoffs among competing constraints. We find that different remodeling strategies produce distinct outcomes. Global gradient descent efficiently drives elongation but yields homogeneous, mechanically fragile tissues with high edge susceptibility. In contrast, the local rule LO generates heterogeneous, “robust-by-design” tension landscapes that buffer perturbations. This connection opens a pathway to classify morphogenetic strategies by their different tradeoffs among competing possible biological priorities, providing a unifying language to describe the importance of energy efficiency, robustness, and evolutionary constraints in developmental systems.

While our initial work focuses on mechanical edge susceptibilities as a measure of robustness, future studies could also directly investigate the robustness of different tuning strategies to specific time-varying perturbations, such as laser ablation [9] or localized optogenetic perturbations to myosin [23]. One could also investigate environmental perturbations such as temperature changes or exposure to environmental toxins. An interesting question that emerges is whether the local rule LO is generically more robust to all of these types of perturbations compared to gradient descent, and whether this might be related to configurational entropy of the states [60].

A related set of questions involves the choice of cost function for the learning rule. Here, we defined the task as pure global shear, which is ultimately related to the concept of ‘developmental checkpoints’ [61, 62] – e.g. Drosophila must extend its body axis far enough at this stage in order to move successfully to the next stage of development [63]. In other words, we decided that it was reasonable to start with the hypothesis that the purpose of this developmental process was to sufficiently elongate the body axis. Many developmental biologists and biophysicists heuristically think and talk about morphogenetic processes in this way, and cost functions provide a quantitative method to encode this information and then test hypotheses.

On the other hand, one could envision many other things that may be important for the embryo to accomplish at the same time – perhaps conserve energy, be robust to specific environmental conditions, etc. One path forward would be to develop and study cost functions that encode these tradeoffs directly, for example supplementing Eq. 4 with an additional term that penalizes mechanical energy expenditures. Although the space of possibilities is vast, it should be possible to systematically study the impact of tuning processes and tradeoffs, and to search for signatures of these processes in developmental systems. Different rules and different tradeoffs should also react differently to fast perturbations like laser ablation or optogenetic tools, which could provide a useful avenue for validation.

One of the utilities of the cost function framework we describe here is that it should be immediately extendable to other types of morphogenetic flows. Although we focused on the simplest deformation, global pure shear, it is straightforward to replace the cost function given by Eq. 4 with one that encodes an arbitrary spatial pattern: e.g. a sink at one location, a shear flow in another, an extensile flow in a third location. This would be exciting to study in more complicated morphogenetic processes, and may also be useful for thinking about designing flows in non-biological active materials [64].

Other extensions could involve more realistic simulations of the tissue models or the tuning processes. For example, simulating CE on curved surfaces could test how tension anisotropy interacts with tissue geometry, which is potentially important in ellipsoidal embryos. One could also include additional features in the model – such as cell turnover – or develop 3D models that include apical cell constriction [65–67]. And although we focused here on very simple, non-dynamic local tuning rules, it should be possible to study more complicated rules with internal degrees of freedom that couple tension to mysoin dynamics with time delays. A challenge will be to understand how the time delays in the local rules interact with the learning rates to generate emergent behavior.

By embedding mechanical adaptation within the tunable matter framework, we provide a quantitative platform to reverse-engineer morphogenetic control. Our results suggest that simple mechanical objectives, when paired with local feedback rules, may suffice to generate robust and evolvable developmental strategies.

## ACKNOWLEDGMENTS

This research was supported in part by grants from the NSF (DMS-2235451) and Simons Foundation (MPS-NITMB-00005320) to the NSF-Simons National Institute for Theory and Mathematics in Biology (NITMB). MLM and SA were supported by Simons Foundation #454947 and NSF-DMR-1951921, and MLM acknowledges additional support from grant number 2023-329572 from the Chan Zuckerberg Initiative DAF, an advised fund of Silicon Valley Community Foundation. AJL acknowledges support of the NSF through DMR-MT-2005749 and the Simons Foundation through Investigator grant #327939, and is also grateful for the hospitality of the Aspen Center for Physics (NSF grant PHY-2210452).

## SUPPLEMENTARY INFORMATION

### Energy minimization

All simulations were carried out with the open-source CellGPU package [43]. We initialize each tissue by placing *N* cell centroids uniformly at random in a periodic square domain of side length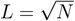, so that the mean cell area is unity. The initial polygonal network is then constructed via a periodic Voronoi tessellation of these centroids, which defines both the vertex positions and cell–cell connectivity.

Each cell *i* is represented as an *n*_*i*_-sided polygon with vertices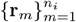 ordered counter-clockwise (and **r**_*n* +1_ ≡ **r**_1_). The cell area and perimeter are

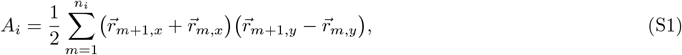

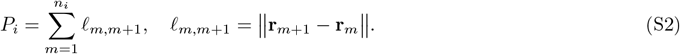

The total tissue energy at fixed tensions *T*_*ij*_ and shear strain *ϵ* Is

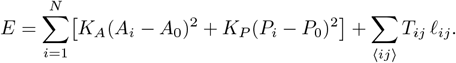

The force on vertex *m* is then

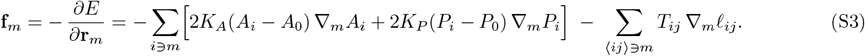

We relax to mechanical equilibrium (force-balanced) at fixed *T*_*ij*_ (and *ϵ* when allowed) using the FIRE algorithm [68]. The stopping criterion is

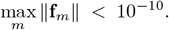

Whenever an edge shrinks below the T1 threshold 𝓁_T1_ = 10^−2^, a neighbor–exchange is performed and FIRE [68] is resumed. In protocols that deform the box via internal tension remodeling (global-GD or local tension rules), the pure-shear strain *ϵ* is treated as an auxiliary variable and updated by enforcing *∂E/∂ϵ* = 0 at each relaxation step.

For dynamics with adaptive tensions, we alternate (1) updating {*T*_*ij*_} according to the chosen feedback rule, and (2) a full FIRE relaxation of {**r**_*m*_} (and *ϵ*) under the new tensions. This two-stage loop ensures the tissue is always in a force-balanced configuration before the next tension update.

### T1 transitions and tension resetting

Whenever an edge *ij* shrinks below the T1 threshold

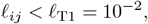

we perform a neighbor-exchange (T1) by flipping that edge: the two vertices are reconnected to their alternate neighbors, creating a new edge of length 𝓁_*ij*_ = 2𝓁_T1_ (see Fig. S1). To prevent repeated back-and-forth flips on the same pair of vertices [69, 70], we reset the tension on the new edge to the tissue-average tension:

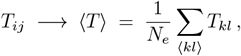

where *N*_*e*_ is the total number of edges. Immediately after resetting *T*_*ij*_, we resume the FIRE relaxation of all vertex positions (and box shear *ϵ*, if present) under the updated tension field, until the force-balance criterion max_*m*_ ∥**f**_*m*_∥ *<* 10^−10^ is again met.

**FIG. S1.**
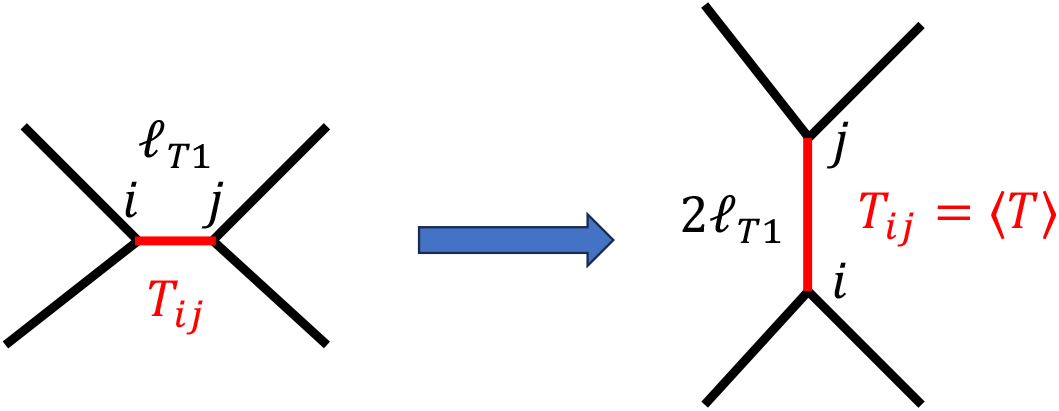
T1 transition and tension resetting. A T1 transition is triggered manually when a cell–cell edge shrinks below a specified cutoff length. As illustrated schematically, in the new configuration, the newly formed edge is initialized by slightly separating its vertices (twice the cutoff length), and its tension is reset to the average tension across all edges in the tissue.

### Computing the shape alignment index *Q*

The cell shape alignment parameter *Q* quantifies both the degree of cellular shape anisotropy and the coherence of orientation within epithelial tissues [51]. Unlike conventional nematic order parameters that measure only directional alignment, *Q* incorporates information about both the elongation of individual cells and the consistency of their orientation across the tissue.

A value of *Q* ≈ 0 indicates either that cells are nearly isotropic (circular) or that elongated cells lack orientational coherence. Values of *Q >* 0 signify that cells are anisotropic (elongated) and that their elongation axes exhibit directional alignment. This parameter is mechanistically significant because it reflects the presence of anisotropic stresses within tissues. Tissues with high *Q* often exhibit direction-dependent mechanical responses and facilitate anisotropic deformation during morphogenetic processes. In the following section, we describe step-by-step how this parameter is computed using a triangulation-based approach [58, 71, 72].

#### Triangulation of Cellular Network

For a given tissue configuration:

1. Compute cell barycenters (geometric centers): **r**_*α*_ = (*x*_*α*_, *y*_*α*_) for each cell *α*
2. Generate triangles from cellular vertices: For each vertex where exactly 3 cells meet, create a triangle connecting their barycenters. (see Fig. S2 a,b)

#### Triangle Shape Tensor

For each triangle *m* with vertices **r**^*A*^, **r**^*B*^, **r**^*C*^ (ordered counter-clockwise):

1. Construct the deformation matrix:

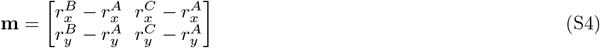
2. Compute shape tensor relative to reference equilateral triangle:

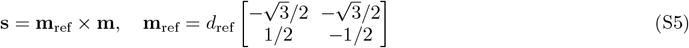

Where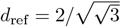 scales to unit area.

#### Triangle Elongation Tensor

1. Decompose **s** into isotropic, deviatoric, and antisymmetric parts:

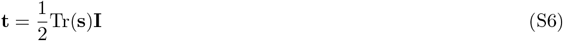

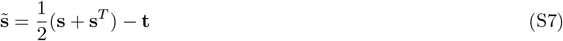

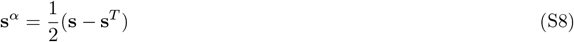
2. Compute rotation angle *θ*:

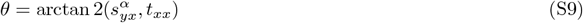
3. Calculate magnitude of deviatoric component:

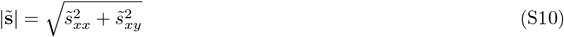
4. Compute elongation tensor **q**:

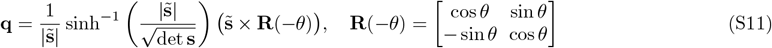

The tissue-level alignment tensor is the area-weighted average:

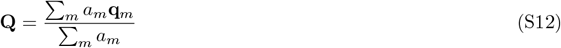

where triangle area *a*_*m*_ is computed via shoelace formula.

#### Scalar Alignment Parameter

The scalar *Q* is the norm of the symmetric traceless tensor:

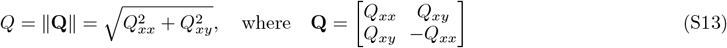

Figure S2 c shows the scalar alignment parameter *Q* as a function of strain for different methods. In the externally driven shear case, *Q* increases monotonically with strain, reflecting the expected alignment of cells as the tissue is stretched. In contrast, the global gradient-descent (GD) method exhibits little change in *Q*, consistent with the strongly suppressed cell-shape anisotropy characteristic of this protocol. The length- and orientation-dependent local rule, however, produces a clear increase in *Q* with strain, confirming that cells become progressively more aligned as the tissue remodels according to this local feedback mechanism.

### Different orientation-dependent local rules

Figure S3 compares the effects of three distinct orientation-dependent local rules that modulate junctional tension: a step function, a triangular function, and a cosine-based rule, cos(2*θ*). Despite their different functional forms, all three produce comparable average cell shape–strain relationships, indicating that the macroscopic behavior is robust to the specific choice of this orientation-dependent local rule.

**FIG. S2.**
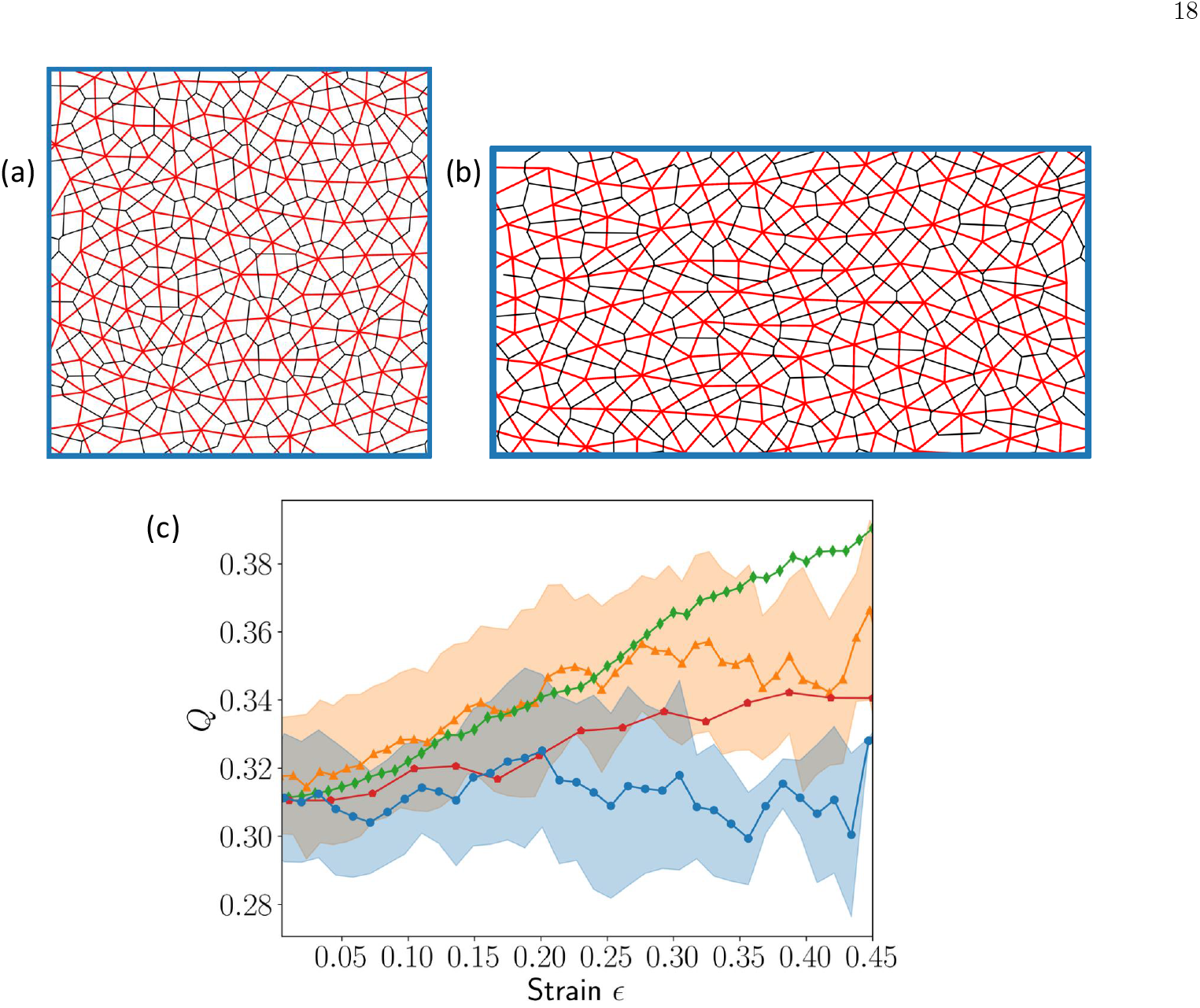
Triangulation of tissues used to compute the alignment parameter *Q*. (a) Triangulated network at low shear stra (b) Triangulated network at higher strain, both corresponding to simulations using the local tension rule 𝓁 cos(2*θ*). (c) Scalar alignment parameter *Q* as a function of strain for different methods: external shear (green diamonds), local 𝓁 cos(2*θ*) r (orange triangles), local cos(2*θ*) rule (red pentagons), and global gradient descent (blue circles).

**FIG. S3.**
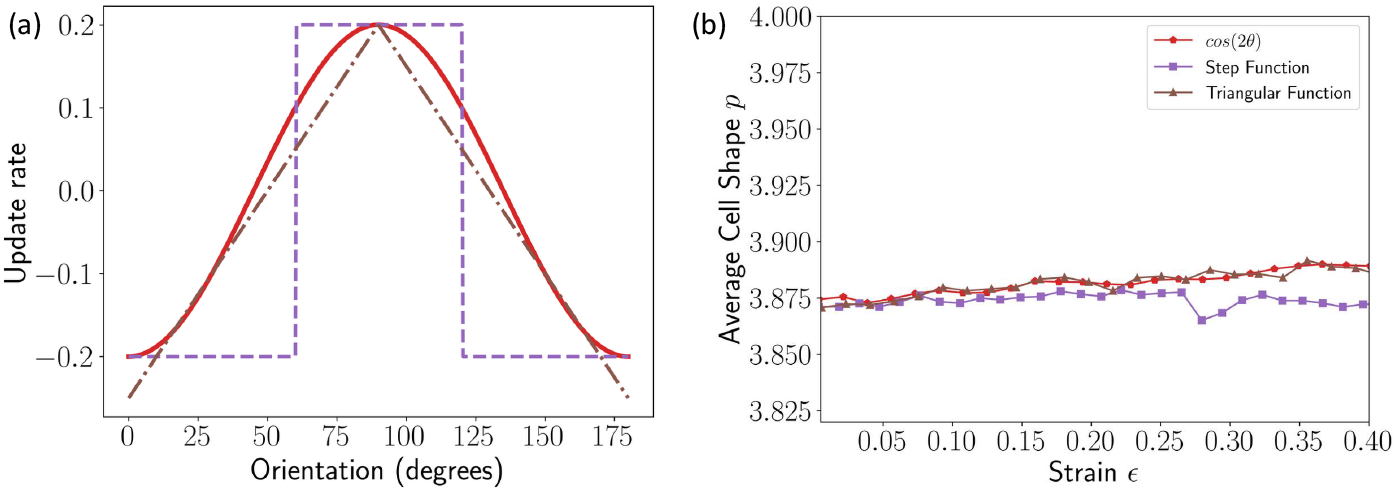
Effect of different orientation-dependent local rules on tissue behavior. (a) Functional forms of the three rules: cos(2*θ*) (solid red), step function (dashed purple), and triangular function (dash-dotted brown). (b) Average cell shape versus strain for each rule, showing that the overall response is qualitatively similar across these rules.

### Computing the edge susceptibility magnitude *s*_*i*_

To probe how individual edges respond to mechanical perturbations, we utilize the concept of *edge susceptibility*, following the approach in Guzmán et al. [59]. Edge susceptibility quantifies how sensitively the length of a given edge changes in response to infinitesimal forces applied at tissue vertices. It provides a local measure of mechanical softness or responsiveness, capturing how edge geometry couples to global force distributions.

Formally, for each edge *i*, we define a susceptibility vector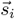, whose components describe how the length 𝓁_*i*_ of edge *i* changes in response to a unit force applied at each vertex *k* in the tissue. For a system with *N*_*v*_ vertices, the susceptibility vector is:

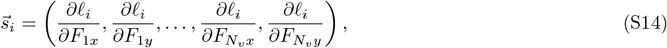

where *F*_*kx*_ and *F*_*ky*_ denote force components applied at vertex *k* in spatial dimensions *x* and *y*. This definition reflects the linear response of edge *i* to perturbations of *physical degrees of freedom* (vertex positions), analogous to voltage responses to node currents in resistor networks [59].

To summarize this information in a scalar form, we compute the squared Euclidean norm of the susceptibility vector:

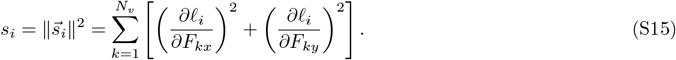

This scalar measure *s*_*i*_ quantifies the *overall mechanical sensitivity* of edge *i*, with larger values indicating edges that undergo significant length changes under force perturbations.

#### Computation via physical Hessian

The susceptibility vector is derived from the inverse physical Hessian **H**^−1^, where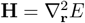 is the Hessian matrix of the energy *E* with respect to vertex positions **r**:

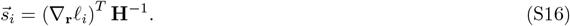

Here, ∇_**r**_𝓁_*i*_ is the gradient of edge length 𝓁_*i*_ with respect to vertex coordinates. This formulation leverages the network’s inherent physical response, independent of adaptive tensions *T*_*j*_.

Figure S4 illustrates the spatial distribution of the components of the edge susceptibility vector for a representative edge. Vertices located near the selected edge exhibit larger values of *s*_*i,α*_, whereas distant vertices show smaller responses. This pattern reflects the expected localized mechanical response of the network to force perturbations. Analyzing the susceptibilities of all edges across the tissue provides insight into the system’s fragility under perturbations and offers a way to assess the robustness of solutions generated by different dynamical methods.

### The projection of local rules onto the global gradient descent

To quantify the similarity between the global gradient descent (GD) rule and the local rules, we computed the projection of the tension update vectors generated by each local rule onto the GD direction (Fig. S5). A positive projection indicates that the local update aligns, at least partially, with the GD update. This alignment explains why both local rules are capable of driving the system toward solutions, despite not being strictly identical to the GD rule. We find that the projection is consistently positive across strain values. Moreover, the cos(2*θ*) rule exhibits a stronger projection onto GD compared to the 𝓁 cos(2*θ*) rule, indicating a closer alignment. This observation is consistent with our analysis of the resulting solutions: the cos(2*θ*)-based solutions show greater similarity to GD solutions than those obtained with the 𝓁 cos(2*θ*) rule.

**FIG. S4.**
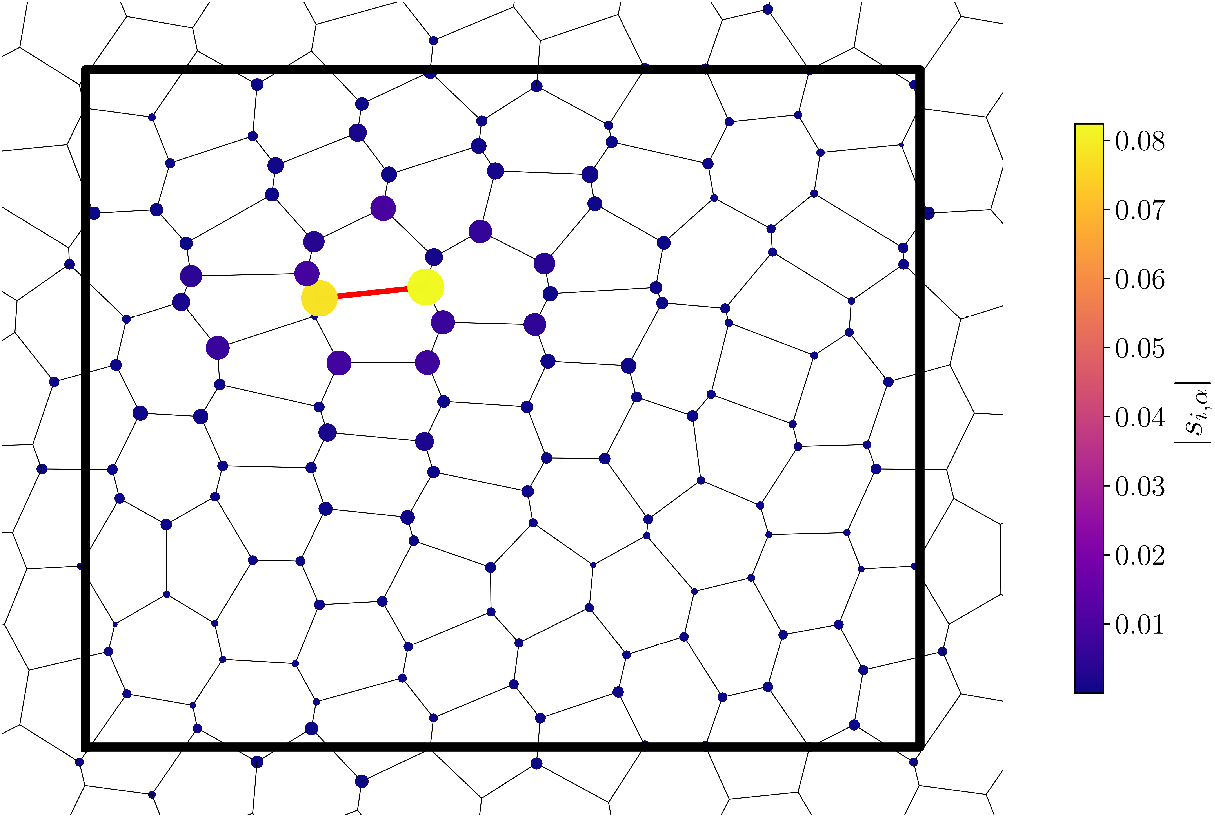
Components of the edge susceptibility vector. For the highlighted edge *i* (red), the *x*-components of the susceptibility *s*_*i,x*_ are shown on all vertices. Vertex size is proportional to the magnitude of *s*_*i,x*_, and vertex color corresponds to the color scale shown on the right.

### Fitting the distributions

We analyzed the statistical properties of (i) total edge tension, (ii) edge susceptibility magnitudes, and (iii) the difference between maximum and minimum edge tension per cell, at a representative strain of *ϵ* = 0.4, by fitting their distributions to gamma probability density functions (Fig. S6).

The gamma distribution with shape parameter *α >* 0 and rate parameter *β >* 0 is defined as

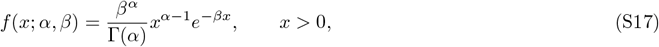

where Γ(*α*) is the gamma function. The mean and variance are given by

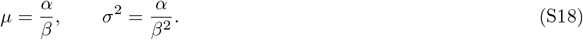

Here, *α* controls the shape of the distribution: for small *α* the distribution is skewed and heavy-tailed, while larger *α* leads to distributions that are more symmetric and Gaussian-like. The parameter *β* acts as the rate (inverse scale), controlling the width: larger *β* corresponds to a narrower distribution concentrated near the mean, while smaller *β* corresponds to a broader, heavier-tailed distribution.

Across all rules and observables, we find that the gamma distribution provides an excellent fit, with coefficients of determination *R*^2^ *>* 0.95. This confirms that the gamma family is well-suited to capture the statistical variability of microscopic observables in our model.

The fitted gamma parameters further clarify systematic differences between global and local rules. For the total tension distributions, the global gradient descent (GD) method exhibits a larger *β* parameter than the local 𝓁 cos 2*θ* rule, consistent with a narrower distribution. In contrast, the local rule yields a broader distribution with heavier tails, reflecting the presence of edges with anomalously high tensions. For the susceptibility distributions, GD shows a smaller *β* compared to the local rule, consistent with broader susceptibility fluctuations under GD. Finally, for the maximum–minimum edge tension difference per cell, GD exhibits a much larger *β* than the local rule, signifying a tighter distribution of tension heterogeneity across cell interfaces.

**FIG. S5.**
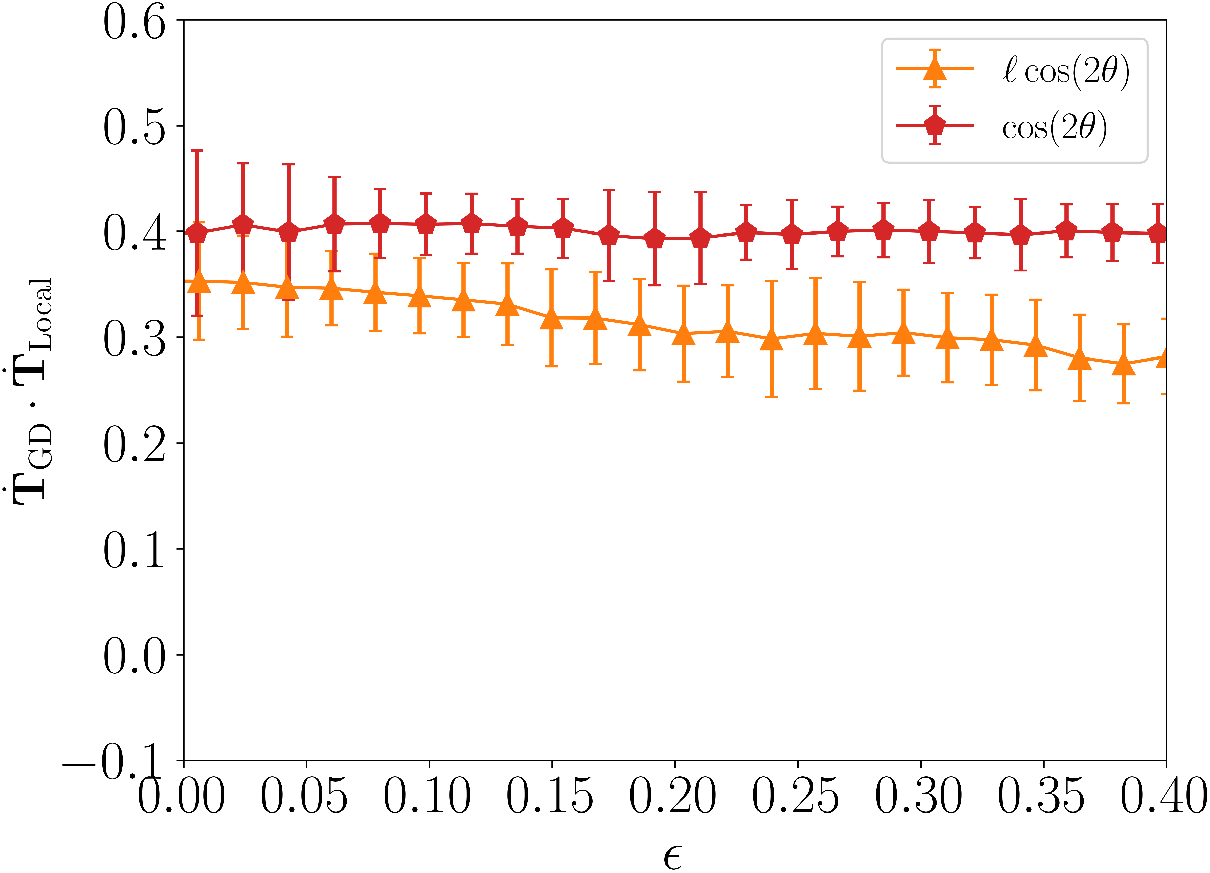
Projection of local rules onto the global gradient descent (GD) update direction. The projection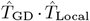 is shown as a function of strain *ϵ* for the 𝓁 cos(2*θ*) rule (orange triangles) and the cos(2*θ*) rule (red pentagons). Both projections remain positive, indicating that the local rules are at least partially aligned with the GD direction. The cos(2*θ*) rule is more strongly aligned with GD than the 𝓁 cos(2*θ*) rule, consistent with the closer similarity of its solutions to GD-based solutions.

### Results for active tension network model

In the standard vertex model we discussed above, edge tensions can have both passive and active contributions. The passive contribution arises indirectly from the perimeter elasticity term: cells with a target perimeter *P*_0_ resist deformation of their perimeter, and this resistance manifests as an effective edge tension even in the absence of any explicitly defined tension variables.

To disentangle these contributions and focus exclusively on active interfacial mechanics, we turn to the Active Tension Network model introduced by Noll et al. [44]. In this model, the perimeter elasticity is removed entirely, and edge tensions *T*_*ij*_ are treated as independent dynamical variables that evolve according to prescribed rules.

The mechanical energy in the ATN model is defined as:

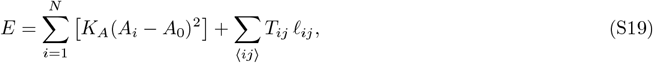

where *A*_*i*_ is the area of cell *i* and *T*_*ij*_ is the tension along the edge shared between neighboring cells *i* and *j*. Eliminating the nonlinear perimeter elasticity makes this model numerically unstable and computationally demanding [50], making it difficult to obtain a large ensemble of converged solutions. For the solutions that were successfully obtained, the overall relationship between cell shape and strain remains consistent with the results from the model that includes perimeter elasticity (Fig. S7).

**FIG. S6.**
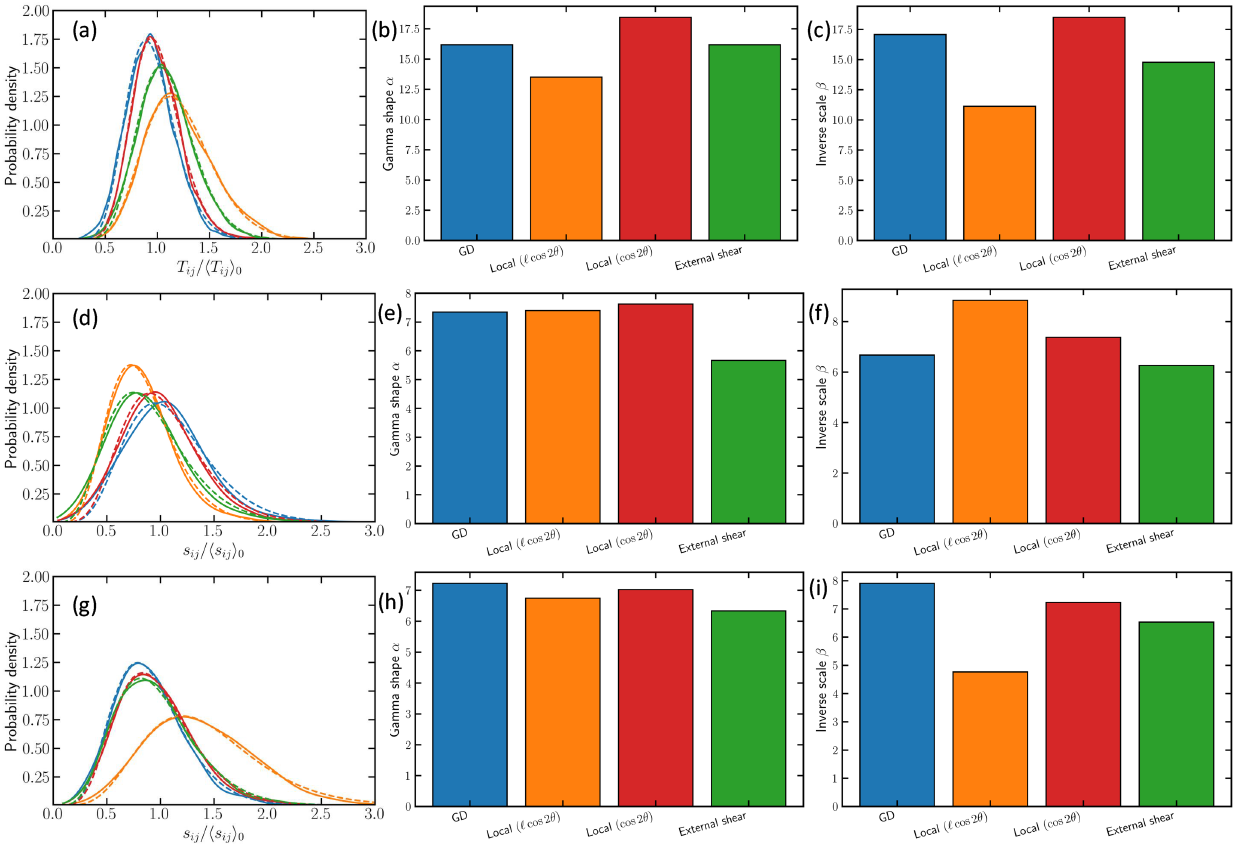
Gamma distribution fits for the distributions of (a–c) total tension, (d–f) edge susceptibility magnitudes, and (g–i) difference between maximum and minimum edge tensions per cell at strain *ϵ* = 0.4, for different methods. Solid lines show the data distributions, dashed lines are gamma distribution fits, and extra panels on the right summarize the fitted gamma parameters *α* and *β* = 1*/*scale for each case. All fits yield *R*^2^ *>* 0.95, demonstrating the suitability of the gamma distribution.

**FIG. S7.**
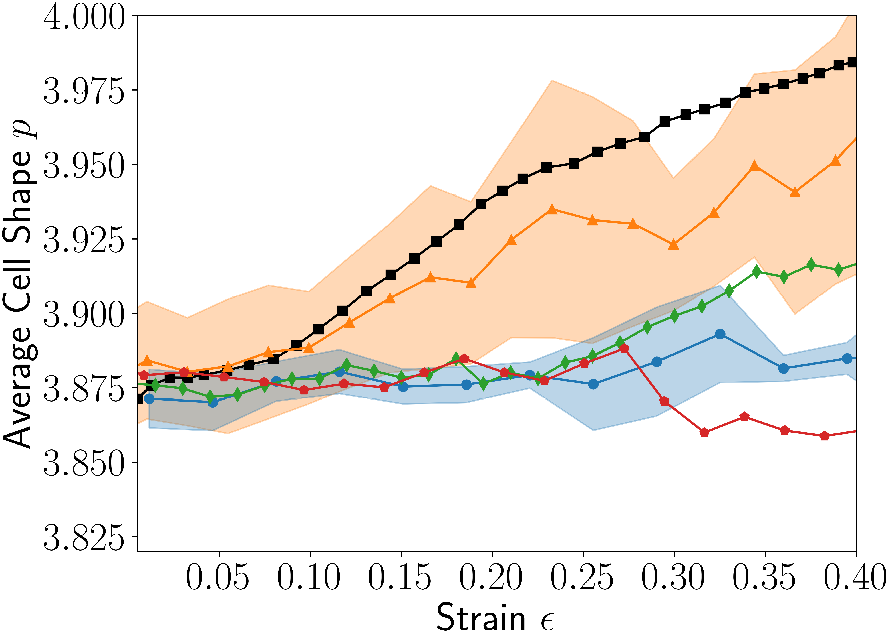
Average cell shape as a function of strain for the active tension network model defined in (S19). Results are shown for different methods: local 𝓁 cos(2*θ*) rule (orange triangles), external shear (green diamonds), local cos(2*θ*) rule (red pentagons), and global gradient descent (blue circles). Experimental data from Ref. [51] are shown as dark squares for comparison.

